# Molecular epidemiology of third-generation cephalosporin-resistant Enterobacteriaceae from Southeast Queensland, Australia

**DOI:** 10.1101/2020.02.20.958728

**Authors:** AG Stewart, EP Price, K Schabacker, M Birikmen, PNA Harris, K Choong, S Subedi, DS Sarovich

**Affiliations:** Department of Infectious Diseases, Sunshine Coast University Hospital, Birtinya, QLD, Australia; Centre for Clinical Research, University of Queensland, Herston, QLD, Australia; GeneCology Research Centre, University of the Sunshine Coast, Sippy Downs, QLD, Australia; Sunshine Coast Health Institute, Birtinya, QLD, Australia; Pathology Queensland, Central Microbiology, Royal Brisbane and Women’s Hospital, Herston, QLD, Australia; Department of Microbiology, Pathology Queensland, Sunshine Coast University Hospital, Birtinya, QLD Australia

**Keywords:** Antibiotic, antimicrobial, resistance, cephalosporin, ESBL, whole-genome sequencing, real-time PCR, Enterobacteriaceae

## Abstract

Third-generation cephalosporin-resistant (3GC-R) Enterobacteriaceae represent a major threat to human health. Here, we captured 288 3GC-R Enterobacteriaceae clinical isolates from 264 patients presenting at a regional Australian hospital over a 14-month period. Alongside routine mass spectrometry speciation and antibiotic sensitivity testing, isolates were examined using rapid (∼40 min) real-time PCR assays targeting the most common extended spectrum β-lactamases (ESBLs; CTX-M-1 and CTX-M-9 groups, plus TEM, SHV, and an internal 16S ribosomal DNA control). AmpC CMY β-lactamase prevalence was also examined. *Escherichia coli* (80.2%) and *Klebsiella pneumoniae* (17.0%) were dominant, with *Klebsiella oxytoca, Klebsiella aerogenes* and *Enterobacter cloacae* infrequently identified. Ceftriaxone and cefoxitin resistance were identified in 97.0% and 24.5% of *E*. coli and *K. pneumoniae isolates*, respectively. Consistent with global findings in Enterobacteriaceae, most (98.3%) isolates harbored at least one β-lactamase gene, with 144 (50%) encoding *bla*_CTX-M-1_ group, 92 (31.9%) *bla*_CTX-M-9_ group, 48 (16.7%) *bla*_SHV_, 133 (46.2%) *bla*_TEM_, and 34 (11.8%) *bla*_CMY_. A subset of isolates (n=98) were subjected to whole-genome sequencing (WGS) to identify the presence of cryptic resistance determinants, and to verify genotyping accuracy. WGS of β-lactamase negative or carbapenem-resistant isolates identified uncommon ESBLs and carbapenemases, including *bla*_NDM_ and *bla*_IMP_, and confirmed all PCR-positive genotypes. We demonstrate that our PCR assays enable the rapid and cost-effective identification of ESBLs in the hospital setting, which has important infection control and therapeutic implications.

## Introduction

Third-generation cephalosporin-resistant (3GC-R) Enterobacteriaceae represent a significant threat to human health due to their ability to rapidly transmit antimicrobial resistance (AMR) determinants within and among bacterial populations (1). Both infection and colonization with these globally-disseminated organisms are associated with poor clinical outcomes and death (2). 3GC-R Enterobacteriaceae prevalence rates vary considerably among hospitals and countries, with particularly high rates in India, Asia, and the Middle East (3). Australia has historically observed a relatively low frequency of 3GC-R Enterobacteriaceae. However, this pattern may be changing, with an extended spectrum β-lactamase (ESBL) phenotype detected in 13.3% of *E. coli* and 9.8% of K. *pneumoniae* in blood culture isolates in 2018 (4), up from 2013 rates of 7.6% and 6.3%, respectively. Moreover, ceftriaxone resistance was seen in 13.4% of *E. coli* and 9.4% of K. *pneumoniae*, with 86.3% and 82.6% containing CTX-M-type ESBL genes, respectively (4).

AMR in 3GC-R Enterobacteriaceae is generally conferred by the presence of an ESBL or the over-expression of a chromosomal or plasmid-borne AmpC β-lactamase (5). CTX-M-type ESBLs are currently the most common AMR determinants found in 3GC-R Enterobacteriaceae isolated in Australian hospitals, with *bla*_CTX-M-1_ and *bla*_CTX-M-9_ groups collectively comprising 86.2% of identified CTX-M-type enzymes (4). Detection of ESBL subtypes is not standard clinical practice; instead, their presence is typically inferred from antimicrobial susceptibility patterns and phenotypic confirmatory tests, which are laborious, costly (∼$AUD25/isolate), and time-consuming (>24h) to perform (6). Use of molecular-based diagnostics for AMR determinant detection in the clinical setting would improve patient outcomes by enabling more rapid, directed, and personalized treatment strategies, thereby improving antibiotic stewardship measures and reducing AMR burden. Their utility has already been demonstrated in a number of clinical settings, including for bloodstream infections, pneumonia, and infection control screening (7–9).

In the current study, we prospectively collected all 3GC-R Enterobacteriaceae identified over a 14-month period at the Sunshine Coast University Hospital, Queensland, Australia, a 450-bed, regional teaching and tertiary hospital. Using large-scale comparative genomics, we developed a novel quadriplex and two singleplex real-time PCR assays to rapidly detect the most commonly identified β-lactamase (including ESBL) genes in 3GC-R Enterobacteriaceae. In association with WGS, we used our diagnostic real-time assays to determine the molecular epidemiology of the β-lactamase genes in our isolate set, and compared their performance with both genotypic and phenotypic typing methods to validate assay accuracy, sensitivity and specificity. Finally, we examined the population structure of our 3GC-R Enterobacteriaceae population to identify potential cases of nosocomial spread or outbreak clusters.

## Materials and Methods

### Ethics

This study was approved by The Prince Charles Hospital Human Resources Ethics Committee (HREC/18/QPCH/110). A waiver of informed consent was granted due to the low to negligible risk associated with this study.

### Isolate collection

Clinical specimens from 264 adult and pediatric patients (including ICU patients [*n=48*]) that were culture-positive for 3GC-R Enterobacteriaceae were collected prospectively from July 2017 through September 2018 at the Sunshine Coast University Hospital microbiology laboratory (*n=*288). Multiple isolates were collected from 23 patients, either due to re-infection over the course of the study (*n=*9), multiple samples taken during initial admission (*n=*11), or collected during the course of their hospital stay (*n=*3). Samples were inoculated onto ChromID™ ESBL agar (bioMérieux) and ChromID™ Carba/OXA agar (bioMérieux) and examined for colored colonies at 18-24 hours. An ESBL screening disc test was performed on isolates growing on ESBL agar. Isolates were speciated using the VITEK® 2 MALDI-TOF MS with the GN ID card (bioMérieux, Murrarie, QLD, Australia). Isolate information is detailed in Table S1.

### Susceptibility testing

Antibiotic susceptibility testing and minimum inhibitory concentration (MIC) values were determined using VITEK® 2 or Etest (bioMérieux) according to the manufacturer’s instructions. EUCAST criteria (v.10.0) were used for susceptibility and resistance interpretation.

### Culture DNA preparation

For PCR, crude DNA was extracted from all isolates using 5% chelex-100 resin (Bio-Rad Laboratories, Gladesville, NSW, Australia). For a subset of samples (*n=50*), a pinhead-sized amount of culture collected using a sterile 10 µL pipette tip was placed directly into 4 µL of PCR mastermix to facilitate rapid turnaround time. For WGS, DNA was extracted using the Quick-DNA miniprep kit (Zymo Research, Irvine, CA, USA).

### PCR development and β-lactamase detection

A quadriplex real-time PCR assay was developed to detect the two major CTX-M ESBL groups (*bla*_CTX-M-1_ and *bla*_CTX-M-9_) and *bla*_SHV_ (both ESBL and non-ESBL); a 16S rDNA probe-based assay (10) was included in this quadriplex to verify DNA integrity. Additionally, two singleplex assays were designed to amplify ESBL and non-ESBL *bla*_TEM_ and *bla*_CMY_ β-lactamase genes (Figure 1). To identify conserved regions for PCR assay design, all publicly available genetic sequences for *bla*_CTX-M-1_, *bla*_CTX-M-9_, *bla*_CMY_, *bla*_SHV_, and *bla*_TEM_ were downloaded from the NCBI Nucleotide database (as of March 2019) and aligned using Clustal Omega v1.2.2. Aligned sequences were visualized in Geneious v11.0.4. Candidate primers and probes were identified in Primer Express v3.0 (Applied Biosystems, Foster City, California, USA) or Primer3 v0.4.0 (11) followed by primer dimer and heterodimer assessment using NetPrimer (http://www.premierbiosoft.com/netprimer/).

**Figure 1:**
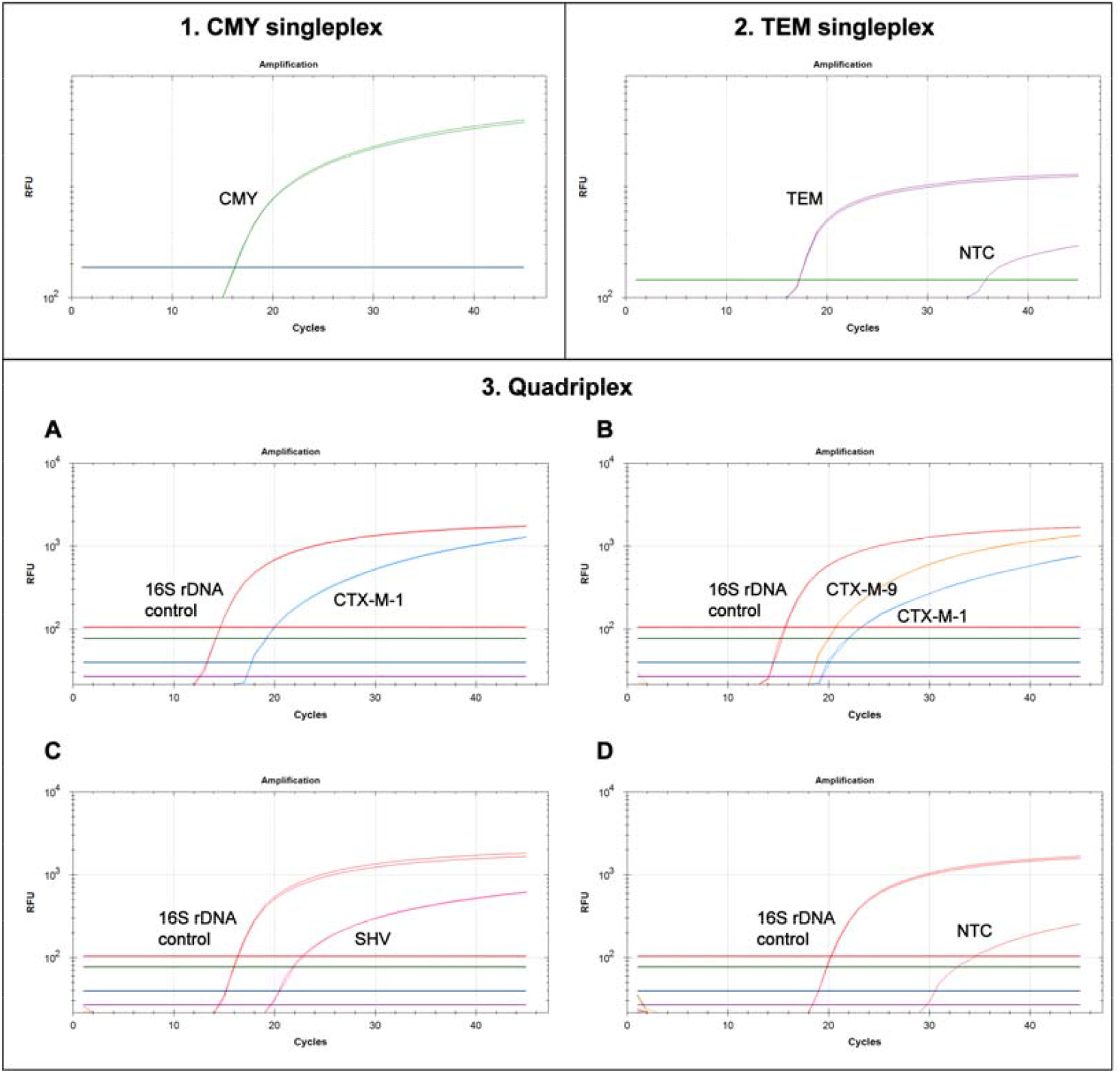
Singleplex and quadriplex PCR performance on third-generation cephalosporin-resistant Enterobacteriaceae. 1: *Enterobacter cloacae* SCHI0016.S.104 containing CMY. 2: *Escherichia coli* SCHI0016.S.205 containing TEM. 3A: *E. coli* SCHI0016.S.187 positive for CTX-M group 1. 3B: *E. coli* SCHI0016.S.60 positive for both CTX-M groups 1 and 9. 3C: E. coli SCHI0046.S.46. 3D: E. coli with only 16s rDNA amplification with no CTX-M or SHV detected. Note that both 16S rDNA and the TEM assays have late amplification in the no-template controls due to low levels of recombinant *E. coli* DNA present in the Sso Advanced Taq polymerase.

Black Hole Quencher probes were used to enable multiplexing in a single PCR well and robust target identification. For the quadriplex assay, the following primers and probes (5’ to 3’; Macrogen Inc., Geumcheon-gu, Seoul, Republic of Korea) were designed: CTX-1 (CTX1_F: CGAMACGTTCCGTCTCGAC, CTX1_R: CTTTCATCCAYGTCACCAGCTG, CTX1_P1: FAM-CGTGATACCACTTCACCTCGGG-BHQ1), CTX-9 (CTX9_F: CGTCSCGCTCATCGATACC, CTX9_R: CAGCTGCTTTTGCGTTTCACTC, CTX9_P1: HEX-CGCTTTCCAATGTGCAGTACC-BHQ1), and SHV (SHV_F: CCARCGTCTGAGCGCC, SHV_R: TATCGGCGATAAACCAGCCC, SHV_P1: Cy5-CAGCACGGAGCGGATCA-BHQ2). The previously described 16S assay (10) incorporated a different fluorophore on the probe (Texas Red-CACGAGCTGACGACARCCATGCA-BHQ2) to accommodate the quadriplex assay format. Two singleplex assays targeting TEM and CMY were also designed: TEM (TEM_F: TGTGGYGCGGTATTATCCCGT, TEM_R: TGCATAATTCTCTTACTGTCAWGCCA, TEM_P1: Cy5.5-CACCAGTCACAGAAAAGCATCT-BHQ2), CMY (CMY_F: TGGCGTATTGGYGATATGTA, CMY_R: TTATGCACCCATGAGGCTTT, CMY_Pr: FAM-TGGGAGATGCTGAACTGGCC-BHQ1). Coincidentally, the reverse primer sequence for our CMY assay was identical to one designed previously (12).

The quadriplex PCR was carried out using 0.1 µM 16S probe and primers, 0.2 µM CTX-1 and CTX-9 probes and primers, 0.3 µM SHV primers, and 0.4 µM SHV probe. For the TEM singleplex, 0.2 µM TEM primers and probe were used, and for the CMY singleplex, 0.2 uM primers and 0.4 probe were used. All reactions contained 1X Sso Advanced Universal Probes Supermix (Bio-Rad), primers, probe/s, 1 µL DNA template, and molecular grade H_2_O, to a 5 µL total volume. PCRs were performed on a CFX96 Touch Real-Time PCR Detection System using white Hard Shell 96-well PCR plates and Microseal ‘C’ Adhesive seals (Bio-Rad) or 0.2 mL white 8-tube PCR strips and optical caps (Bio-Rad). For bacterial cultures, thermocycling conditions comprised an initial 2 min denaturation at 95 °C, followed by 45 cycles of 95 °C for 1 sec and 60°C for 3 sec (total run time=39 min).

### LoD and LoQ determination

Three previously characterized strains (13) that represent the PCR assays developed in this study – MER85 (*K. pneumoniae*), MER110 (*E. coli*), and MER132 (*E. coli*) – were subject to total DNA extraction using the Gram-positive bacteria extraction protocol of the Qiagen DNeasy kit (Qiagen, Chadstone Centre, VIC, Australia). DNA quantities were measured on the Qubit 4 fluorimeter (Thermo Fisher Scientific, Seventeen Mile Rocks, Qld, Australia) using the Qubit dsDNA BR kit, and normalized to 40 ng/µL in molecular grade H_2_O. LoD and LoQ values were determined across 1:10 serial dilutions (range: 40 ng to 0.4 fg) using MER85 (*bla*_TEM_ singleplex), MER110 (*bla*_CMY_ singleplex), and a 50:50 mix of MER85 (*bla*_CTX-M-1_ group, *bla*_SHV_) and MER132 (*bla*_CTX-M-9_ group) for the quadriplex assay. For PCR thermocycling, denaturation and annealing times were increased to 5 sec and 10 sec, respectively, to ensure assay robustness under very low DNA input conditions. Genome equivalents were calculated by assuming *E. coli* and *K*. pneumoniae genome sizes of 4.984 Mbp and 5.638 Mbp, respectively.

### Assay specificity

The quadriplex and singleplex assay amplicons were first assessed for specificity *in silico* using the microbial nucleotide discontiguous BLAST tool and the Complete genomes database (http://blast.ncbi.nlm.nih.gov/; performed March 2019). Next, assay specificity was examined by laboratory testing against a range of both pathogenic and non-pathogenic microbial species found humans. This diversity panel comprised *Achromobacter spp. (n=4), Bordetella bronchiseptica (n=1), Candida albicans (n=1), Cupriavidus sp. (n=2), Enterobacter aerogenes (n=1), Enterobacter cloacae (n=3), Enterococcus faecalis (n=1), Klebsiella pneumoniae (n=2), Klebsiella oxytoca (n=1), Lactobacillus spp. (n=2), Prevotella spp. (n=6), Pseudomonas aeruginosa (n=10), Staphylococcus aureus (n=4), Staphylococcus epidermidis (n=1), Stenotrophomonas maltophilia (n=5), and Veillonella spp. (n=4)*.

### Whole-genome sequencing (WGS)

WGS was performed at the Australian Centre of Ecogenomics (University of Queensland, St Lucia, QLD, Australia) on the NextSeq 500 (Illumina, San Diego, CA, USA) with Nextera DNA Flex libraries (Illumina).

### Genomic analysis

All genomic data were lightly quality-filtered using Trimmomatic v0.39 (14) using parameters described elsewhere (15). WGS data were screened for multiple species using Kraken 2 v2.0.8 (16). Single species were assembled using MGAP v1.1 (https://github.com/dsarov/MGAP---Microbial-Genome-Assembler-Pipeline), whereas WGS data containing multiple species were assembled with metaSPAdes (17). To determine plasmid copy number and identity, plasmids were assembled using the plasmid module of SPAdes(18) with plasmid homology determined using BLAST(19). Assemblies were binned with MaxBin v2.0 (20) and speciated with Kraken 2 v2.0.8 (16). Antibiotic resistance genes were identified using the Resistance Gene Identifier (21). Phylogenomic analysis was performed using SPANDx v3.2.1 (22). Samples identified as containing multiple strains according to the method described by Aziz and colleagues (23) were excluded from phylogenetic analysis (*n=7*); samples able to be split into single species using MaxBin were included in phylogenetic analysis (*n=4*). *E. coli* O157:H7 Sakai (NCBI Assembly ID: ASM886v2) (24) or *K. pneumoniae* HS11286 (NCBI Assembly ID: ASM24018v2) (25) were used as read mapping reference genomes for species-specific analyses. To achieve the highest possible resolution, error-corrected assemblies of strains with matching multilocus sequence types (STs) were used as reference genomes for transmission analyses. ST profiles were determined using SRST2 v0.2.0 (26). Phylogenetic trees were reconstructed using PAUP v4.4.7 (27) and visualized in iTOL v5.5(28).

### WGS data

Raw sequence reads generated in this study are available on the NCBI Sequence Read Archive database under BioProject accession number PRJNA606985.

## Results and Discussion

ESBL enzymes have established themselves as major contributors of AMR in Gram-negative bacteria and are increasing in prevalence, particularly in the community setting. Over 14 months, we collected a total of 288 isolates from all diagnosed cases of 3GC-R Enterobacteriaceae in a regional hospital in Southeast Queensland, Australia. Our patient cohort was predominantly female (64.0%) and aged over 65 (60.9%). ICU patient specimens comprised 54 (18.8%) of the total collection, and were obtained from 48 ICU patients over a 12-month period. Five ICU patients had multiple isolates collected over the course of their stay in the ICU. Three patients isolated single species with identical AMR profiles, two patients isolated multiple species: one with identical AMR profiles and one with differing profiles (Table S1). *E. coli* and *K. pneumoniae* were dominant, comprising 80.2% and 17.0% 3GC-R strains, respectively (Table 1 and S1). The remaining isolates were identified as *E. cloacae* (1.4%; *n=4*), *K. oxytoca* (1.0%; *n=3*), and K. aerogenes (0.3%; n=1). Resistance to ceftriaxone, ceftazidime or cefepime was common and observed in 96.2%, 93.4%, and 86.8% of isolates, respectively. In contrast, cefoxitin resistance was noted in only 81 (28.1%) isolates (Table 1). Meropenem resistance was rare, with only six resistant strains (2.1%) identified. The double disc diffusion method for ESBL phenotypic testing was negative in 32 (11.1%) isolates, indicating a lack of β-lactamase inhibition in the presence clavulanic acid. The remaining 256 (88.9%) isolates were positive for this phenotypic test.

**Table 1:**
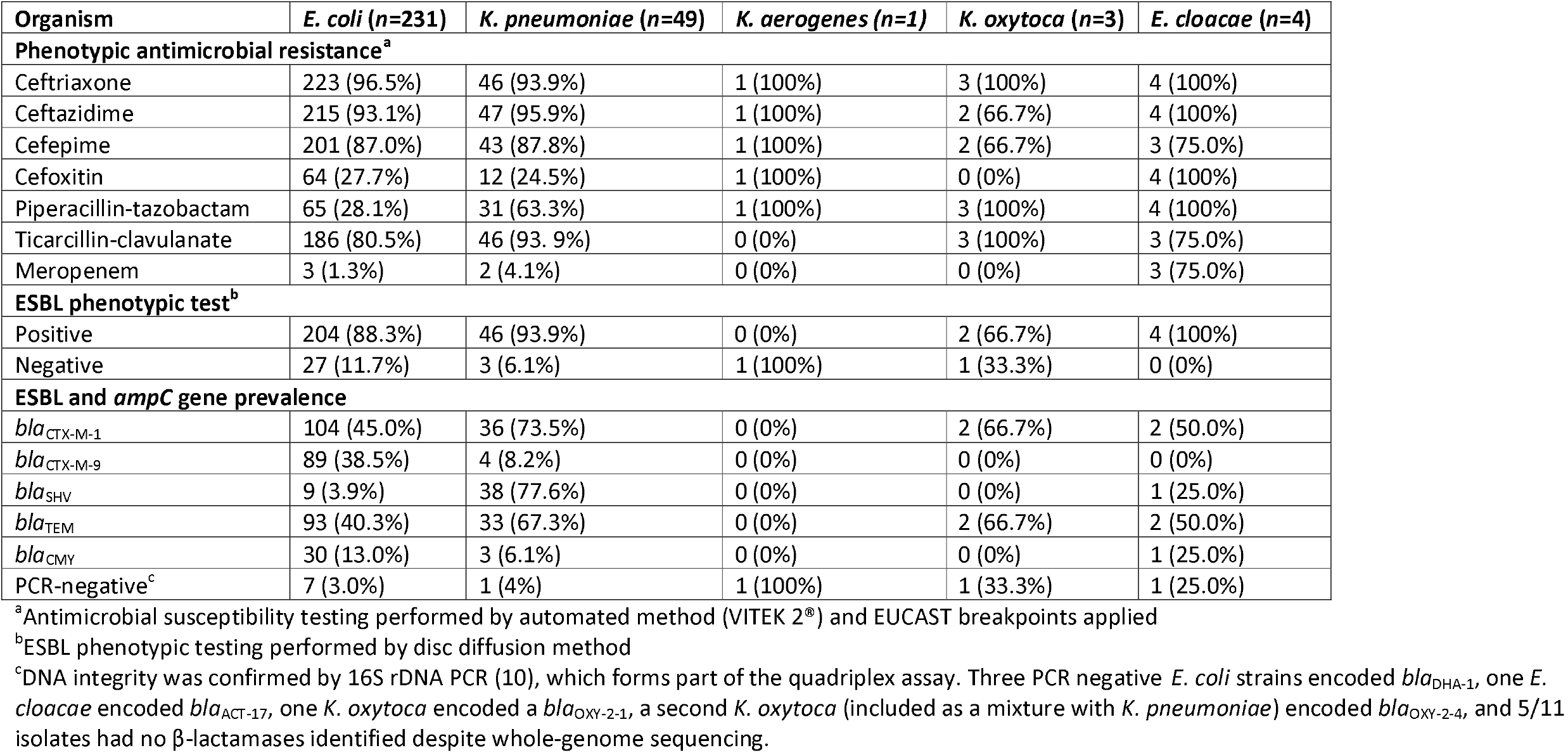
Summary of antimicrobial resistance mechanisms among 288 clinical Enterobacteriaceae isolates retrieved from 264 patients admitted to a Southeast Queensland hospital over a 14-month period.

### ESBL and β-lactamase assay performance

Preliminary validation of our singleplex (*bla*_TEM_, *bla*_CMY_) and quadriplex (*bla*_CTX-M-1_ group, *bla*_CTX-M-9_ group, *bla*_SHV_, 16S rDNA) PCR assays was performed to determine their accuracy and specificity. Following *in silico* verification, assays were tested on 12 previously characterized control strains (Table S2) (29) that represented a range of Enterobacteriaceae containing the gene targets of interest. All expected targets were identified in this control dataset; no false positives or false negatives were detected, although both the *bla*_TEM_ and 16S rDNA assays yielded late (>30 cycles) amplification in no-template control wells (Figure 1). This result was expected due to residual *E. coli* DNA presence in the recombinant Sso Advanced Taq polymerase, which encodes both *bla*_TEM_ and 16S rRNA genes.

We found that all genotypes could be accurately determined for the control strains by directly inoculating PCR wells with a very small amount of bacterial culture, thereby obviating the need for DNA extraction and leading to faster time-to-diagnosis. However, this technique requires skill to ensure that the PCR is not inhibited by excess DNA and cellular constituents, and can lead to greater PCR failures if not performed correctly. In addition, we recommend using 10 µL rather than 5 µL total PCR volume as, in our hands, the lower reaction volume led to a greater failure rate. Inclusion of the 16S rDNA assay enables true negatives to be differentiated from PCR failures. This method provides an attractive option where rapid diagnosis is paramount, or where DNA extraction is not feasible.

Next, we screened our PCR assays across a diversity panel of non-Enterobacteriaceae human pathogens or colonizers (*n=*48 strains). Our assays showed 100% sensitivity and specificity for all targets, with no amplification detected in these isolates, except for the 16S target in bacterial strains; as with the control dataset, the *bla*_TEM_ PCR was inhibited by the presence of non-*bla*_TEM_-encoding DNA and therefore did not amplify in any diversity panel strains.

### LoQ and LoD determination of the ESBL and β-lactamase assays

Using serial dilutions of previously characterized DNA samples ranging from 40 ng to 0.4 fg, the LoQ values for each assay were identified (Figure 2) as follows: *bla*_CMY_, 400 fg (∼73 genome equivalents [GEs]); *bla*_TEM_, 400 fg (∼65 GEs); 16S rDNA, 40 fg (∼3 GEs); *bla*_CTX-M-1_ group, 4 pg (∼344 GEs); *bla*_CTX-M-9_ group, 4 pg (∼344 GEs); and *bla*_SHV_, 4 pg (∼344 GEs). LoD values for each assay were: *bla*_CMY_, 4 fg (∼7 GEs); *bla*_CTX-M-1_ group, 4 pg (∼344 GEs); *bla*_CTX-M-9_ group, 400 fg (∼34 GEs); and *bla*_SHV_, 400 fg (∼34 GEs). Due to low levels of recombinant *E. coli* DNA present in the Sso Advanced Taq polymerase, which results in late amplification in no-template control wells, LoD values for the *bla*_TEM_ and 16S rDNA assays could not be determined. As expected, these results indicate inferior performance of the quadriplex assay when compared with the singleplex assays. Although not examined in this study, increased LoQ and LoD values may be achieved for the *bla*_CTX-M-1_ group, *bla*_CTX-M-9_ group, and *bla*_SHV_ assays by omitting the 16 rDNA assay, or by running these assays in singleplex format. Although we have defined the limits of our assays using pure bacterial templates, it remains to be determined what detection limits would be achieved from clinical specimens.

**Figure 2.**
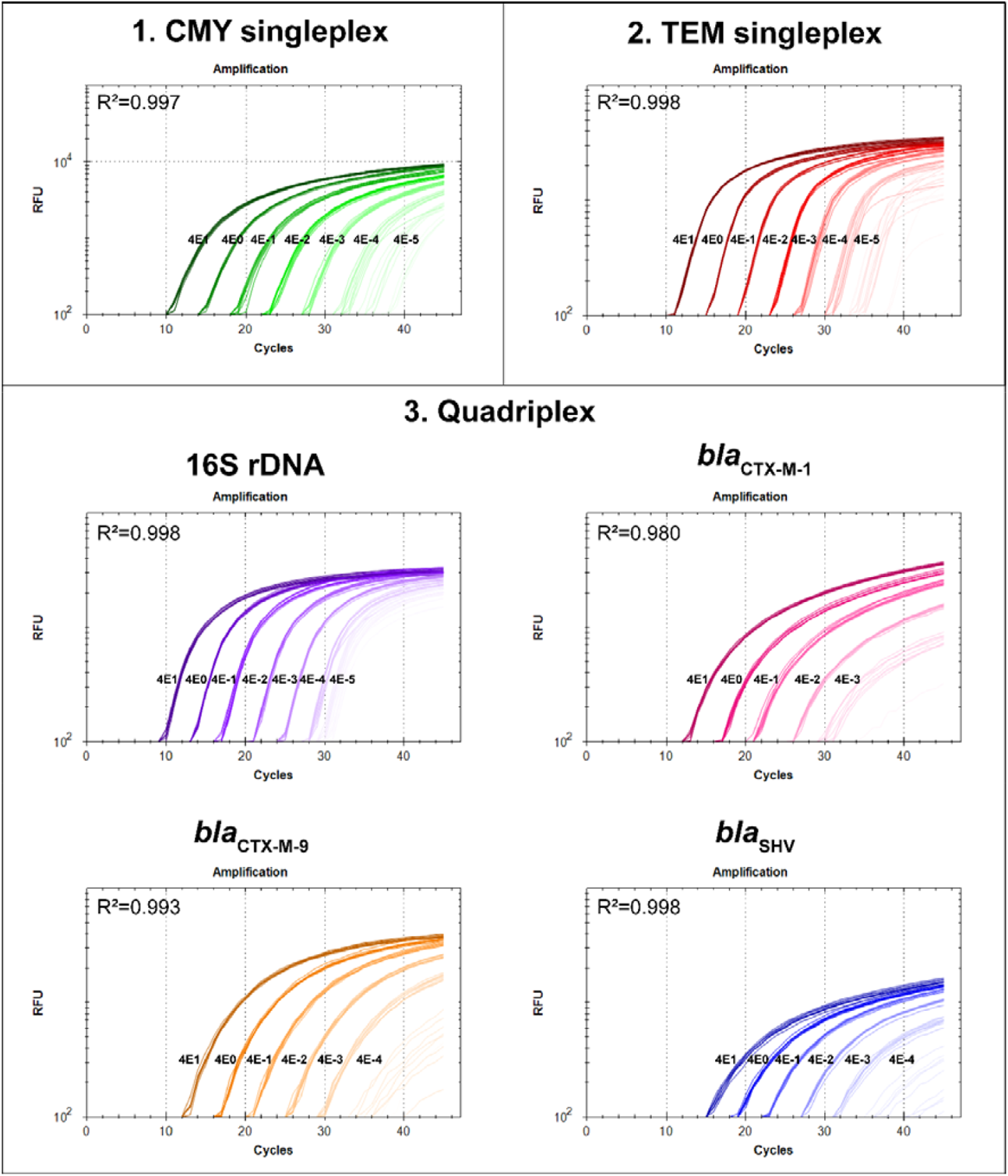
Limits of Detection (LoD) and Quantitation (LoQ) for the extended spectrum β-lactamase real-time PCR assays developed in this study. The *bla*_CMY_ and *bla*_TEM_ assays were examined as singleplex PCRs, whereas the 16S rDNA, *bla*_CTX-M-1_ group, *bla*_CTX-M-9_ group, and *bla*_SHV_ assays were examined in quadriplex format.

### ESBL and β-lactamase prevalence in clinical 3GC-R Enterobacteriaceae isolates

Following specificity confirmation, the PCR assays were tested across our clinical set (*n=*288). All assays performed well, with 100% concordance between PCR genotypes and WGS of 98 isolates, and with no false negatives identified (Table S3). Overall, a β-lactamase genotype was identified in 277 (96.2%) isolates. Consistent with previously reported rates (29), *bla*_CTX-M-1_ group was most common, being identified in 144/288 (50%) isolates, including 105/231 (45.5%) *E. coli* and 36/49 (73.5%) K. pneumoniae. The *bla*_CTX-M-9_ group was next frequent, being found in 93/288 (32.3%) isolates, including 89/231 (38.5%) E. coli and 4/49 (8.2%) *K. pneumoniae*. The presence of both *bla*_CTX-M-1_ group and *bla*_CTX-M-9_ group enzymes was identified in five (1.7%) *E. coli* and one (2.0%) *K. pneumoniae* isolate. Of the eight isolates belonging to other species, *bla*_CTX-M-1_ and *bla*_CTX-M-9_ group genes were present in 4/8 (50%) and 0/8 (0%) of these isolates, respectively. The acquired ampC gene, *bla*_CMY_, was identified in 30/231 (13%) of *E. coli*, 3/49 (6.1%) of K. pneumoniae, and 1/8 (13%) other Enterobacteriaceae species. *bla*_CMY_ and *bla*_CTX-M-1_ were both identified in seven isolates (five *E. coli*, one K. pneumoniae, and one *E. cloacae*) and *bla*_CMY_ and *bla*_CTX-M-9_ were present in two *E. coli* isolates only (Table 1).

Seven strains (five *E. coli and two K. pneumoniae*) had *bla*_SHV_ only; nine *E. coli* had *bla*_TEM_ only, and seven (five K. pneumoniae, one *E. coli*, and one *E. cloacae*) had both *bla*_SHV_ and *bla*_TEM_. Although most SHV (e.g. *bla*_SHV-1_) and TEM (e.g. *bla*_TEM-1_, *bla*_TEM-2_) enzymes do not typically cause 3GC-R, their presence in strains that lack CTX-M-type ESBLs should be investigated, as these enzymes can occasionally confer 3GC-R through enzyme active site mutation or gene overexpression (5). Of the 14 strains encoding *bla*_SHV_ but not *bla*_CTX-M-1_ group, *bla*_CTX-M-9_ group, or *bla*_CMY_ enzymes, five had accompanying WGS data, which permitted assessment of *bla*_SHV_ ESBL status. In all cases, the *bla*_SHV_ variant was an ESBL (two *bla*_SHV-2_, one *bla*_SHV-64_, one *bla*_SHV-66_, and one *bla*_SHV-134_) (30). In contrast, none of the four strains encoding *bla*_TEM_ but not *bla*_CTX-M-1_ group, *bla*_CTX-M-9_ group, *bla*_CMY_, or ESBL-encoding *bla*_SHV_ enzymes were ESBLs, with all being *bla*_TEM-1_. Three of these were negative with the double disc diffusion method, indicating probable chromosomal mutations underpinning this phenotype. The basis for the ESBL phenotype in the fourth strain could not be elucidated.

### Correlation between bla_CMY_ and cefoxitin resistance

Of the 81 cefoxitin-resistant isolates, *bla*_CMY_ was detected in only 31 (38.3%) cases. Although imperfect, cefoxitin resistance is often used as a marker for AmpC presence (32). Correlation is also dependent on the MIC, with higher MICs more reliably predicting AmpC presence (32). Our findings concur with this prior work, with 61.7% of cefoxitin-resistant isolates failing to amplify *bla*_CMY_. AmpC has previously been reported as an underappreciated mechanism of 3GC-R due to a lack of standardized testing (33). As such, molecular methods that can detect other common and emerging β-lactamase gene families (e.g. *bla*_DHA_, *bla*_OXY_, *bla*_MIR_) may prove useful for infection control management and treatment of multidrug-resistant infections. This is particularly true for piperacillin-tazobactam, which may not be effective against AmpC producers (29, 34).

### WGS identifies uncommon ESBL genes

Eleven (3.8%) PCR-negative isolates that otherwise successfully amplified their 16S rDNA internal control (seven *E. coli*, one *K. pneumoniae*, one *K. oxytoca*, one *K. aerogenes* and one *E. cloacae*) were subjected to WGS to determine the molecular basis of 3GC-R. Six were found to carry uncommon β-lactamase enzymes, with three *E. coli* encoding *bla*_DHA-1_, one *K. pneumoniae/K. oxytoca* mixture harboring *bla*_OXY-2-4_, one *K. oxytoca* carrying *bla*_OXY-2-1_, and one *E. cloacae* carrying *bla*_ACT-17_. The remaining five isolates had no discernable 3GC-R determinants, although misclassification via automated susceptibility testing or upregulation of chromosomal β-lactamases are both possible causes for this discrepancy.

Six isolates (three *E. cloacae*, two *E. coli* and one *K. pneumoniae*) exhibiting meropenem resistance alongside 3GC-R were also subjected to WGS to determine the basis of resistance towards this carbapenem. As expected, meropenem resistance was mostly conferred by carbapenemases, with 4/6 (66.6%) isolates possessing one of these enzymes. *bla*_NDM-1_ or *bla*_IMP-4_ were identified in two *E. cloacae* isolates, *bla*_NDM-5_ was identified in an *E. coli* strain, and an OXA-48-like enzyme, *bla*_OXA-232_, was identified in *a K. pneumoniae* isolate. No carbapenemase gene could be identified in the remaining two isolates (one *E. coli* and one *E. cloacae*). Although not investigated in this study, chromosomally-encoded porin loss or efflux pump overexpression may form the basis of meropenem resistance in these strains.

### Monitoring patient-to-patient spread in an ICU

Fifty-four (18.8%) 3GC-R isolates obtained during this study originated from ICU-admitted patients. Due to their dominance, only *E. coli* and *K. pneumoniae* genomes were analyzed for potential transmission events. Among the *E. coli* population (Figure 3), isolates SCHI0016.S.60 and SCHI0016.S.62, and SCHI0016.S.265 and SCHI0016.S.46, were separated by 10 SNPs and 2 SNPs, respectively; however, both pairs came from individual patients, reflecting expected levels of within-host evolution of a single infecting strain. *E. coli* isolates SCHI0016.S.42 and SCHI0016.S.294 were very similar, being separated by 13 SNPs. Although both strains were isolated from ICU-admitted patients, SCHI0016.S.294 was isolated 306 days after SCHI0016.S.42 with no other linking epidemiology identified. Additionally, no similar strains were isolated from ICU patients over this time period suggesting that these do not represent transmission. E. coli isolates SCHI0016.S.172 and SCHI0016.S.274 differed by ∼300 SNPs, with no epidemiological or temporal correlation. ∼300 SNPs separated SCHI0016.S.112 and SCHI0016.S.229, again with no demonstrable epidemiological link. Among *K. pneumoniae* (Figure 4), the closest isolates, SCHI0016.S.15 and SCHI0016.S.261, differed by 361 SNPs, with no epidemiological link. Additionally, no horizontal transfer of plasmids carrying ESBL determinants was observed. As standard practice, all ICU patients, and all patients in other wards infected or colonized with ESBL-producing *E. coli* and *K. pneumoniae*, have single rooms with an assigned nurse (1:1 ratio), with contact precautions exercised. In addition, hand hygiene rates in our hospital ICU are generally above 80% (35). Taken together, our results demonstrate no evidence of transmission in the ICU, suggesting that the implemented hygiene measures are effective.

**Figure 3.**
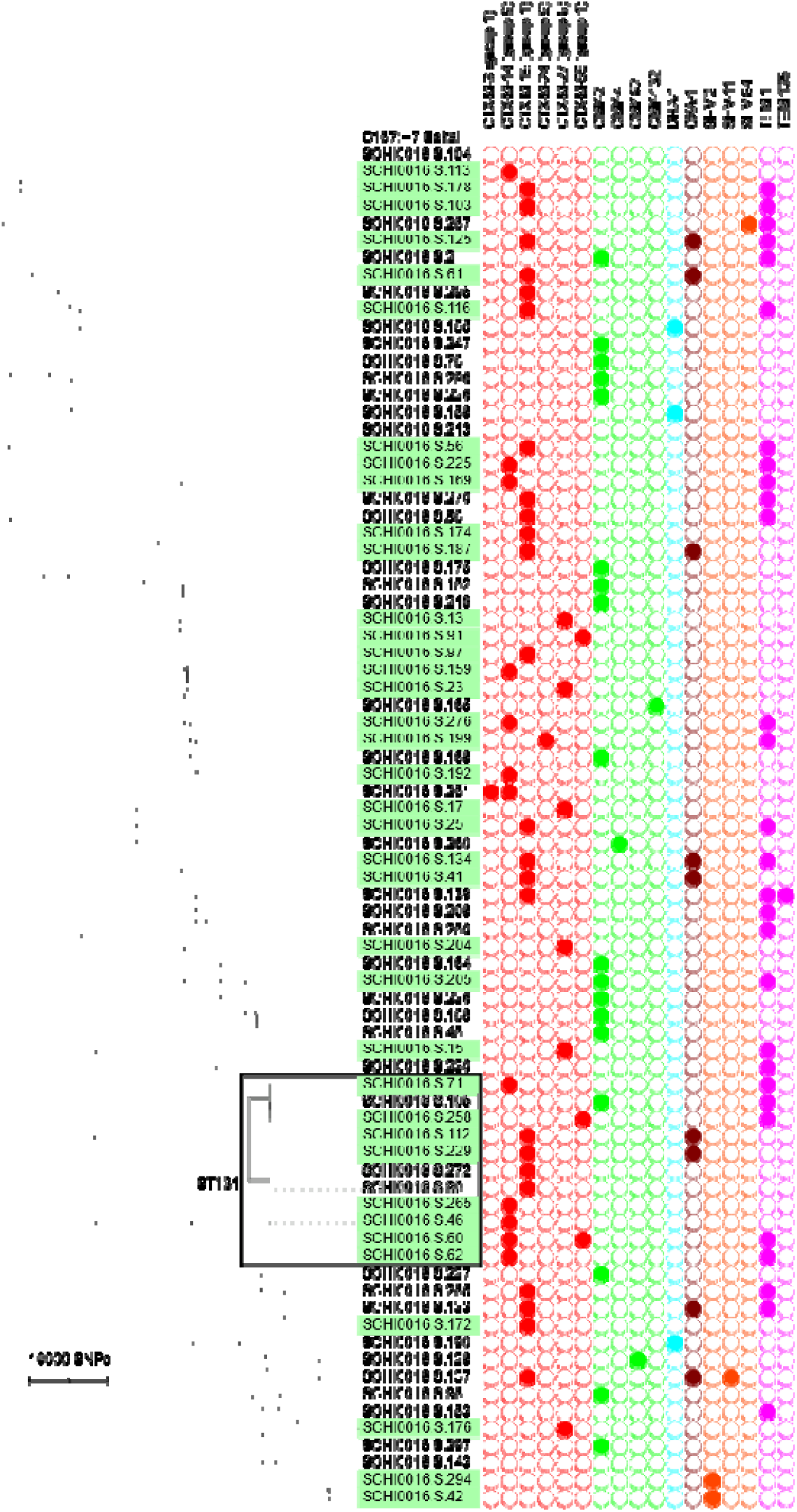
Whole-genome phylogeny of 79 third-generation cephalosporin-resistant Escherichia coli isolates and their β-lactamase gene complement. Sequence type (ST)131 is indicated by the gray-shaded box; strains isolated from intensive care unit patients are highlighted in green.

**Figure 4.**
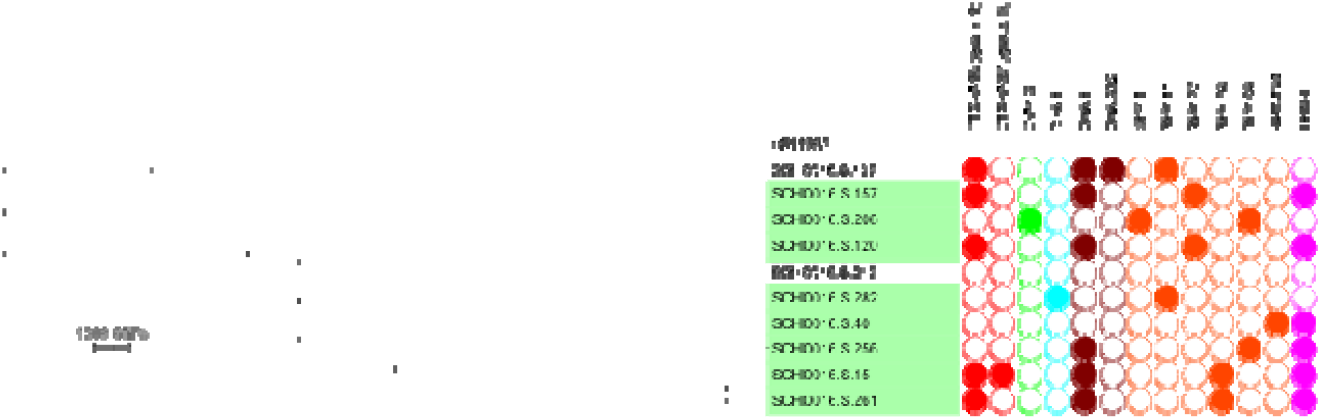
Whole-genome phylogeny and β-lactamase enzyme presence in *Klebsiella pneumoniae*. Green-shaded isolates were isolated from the intensive care unit.

### Population structure of E. coli and K. pneumoniae genomes

We subjected an additional 36 strains to WGS (total n=98) to capture a snapshot of genomic diversity and population structure (Figures 3 and 4). *In silico* MLST of the 79 *E. coli* strains found that ST131 was most common (*n=*11; 13.9%). This finding is consistent with a recent study of the population structure of Australian, New Zealand, and Singaporean multidrug-resistant *E. coli* isolates (29). The highly successful ST131 clone has now spread worldwide, where it frequently causes urinary tract and bloodstream infections and accounts for 12–27% of all community-acquired *E. coli* infections (36, 37). Among our ST131 isolates, we found high frequencies of fluoroquinolone resistance-conferring variants (GyrA_S83L_, GyrA_D87N_, ParC_S80I_ and ParC_E84V_) alongside frequent carriage of plasmid-borne CTX-M-14 (CTX-M-9 group) and CTX-M-15 (CTX-M-1 group) ESBLs. We also identified three strains belonging to the emerging multidrug resistant pandemic clone, ST1193. These isolates also possessed the same three fluoroquinolone-resistant SNPs as ST131. It has been proposed that ST1193 strains evolved fluoroquinolone resistance relatively recently through multiple recombination events with other *E. coli* strains (38).

Phylogenomic analysis of our 10 *K. pneumoniae* genomes identified several distinct lineages, which comprised two ST14 strains and one each of ST17, ST307, ST309, ST395, ST656, ST697, ST873, and ST969. Of note, the globally disseminated ST258 clone, which is responsible for a large number of hospital deaths (39), was not identified in our dataset. However, we identified another globally disseminated clone, ST395, which has also been retrieved from clinical cases in Europe and Asia (40). Concerningly, this ST395 isolate, SCHI0016.S.137, carried both the ESBL genes *bla*_SHV-11_ and *bla*_CTX-M-15_ and the OXA-48-like carbapenemase, *bla*_OXA-232_, the latter of which confers low-level meropenem resistance. Additionally, we observed one *K. pneumoniae* ST307, which is an emerging multi-drug resistant, globally disseminated clone postulated to have emerged in the mid-1990s(41). Although reports of this clone from Australian populations in uncommon it has been previously observed(41).

This study had several recognized limitations. First, isolates were collected from a nosocomial setting; no community sources were sampled. It is well-established that differences in AMR patterns and their associated molecular mechanisms exist between hospital and community settings. Second, the absence of certain high-risk populations within our hospital (e.g. solid organ or hematopoietic stem cell transplantation) limits the generalizability of our findings to other settings. Third, our PCR assays can only identify the presence or change in copy number of the most common ESBL and *ampC* genes. Although these loci make up the majority (>90%) of β-lactamase enzymes present in 3GC-R Enterobacteriaceae, they do not capture all potential ESBLs, leaving less common ESBL and *ampC* genes unidentified. In addition, PCR assays for *bla*_TEM_ or *bla*_SHV_ cannot discriminate between ESBLs and non-ESBLs, with additional assays or concurrent WGS required to confirm ESBL status. Our assays also cannot detect chromosomal changes leading to AMR (e.g. porin mutations or efflux pump up-regulation), making these assays less useful for 3GC-R bacteria that frequently encode these AMR determinants (e.g. *Pseudomonas aeruginosa*). Fourth, our PCR was unable to distinguish chromosomal vs. low-copy plasmid-borne β-lactamase genes, which has implications for AMR transmissibility between and among bacterial populations. Finally, ∼40% of samples were obtained from rectal swabs, which have limitations for detection of multidrug-resistant organisms due to the high microbial burden coupled with difficulties in sample collection. It is therefore likely that we did not capture all 3GC-R isolates during our study period, and thus may have missed potential transmission events.

## Conclusion

Taken together, our findings identified a predominance of CTX-M (and to a lesser extent, CMY) β-lactamases among 3GC-R Enterobacteriaceae in the Sunshine Coast region. These findings are consistent with global findings of Gram-negative bacterial AMR determinants, particularly among *E. coli*. Our study demonstrates the likely absence of 3GC-R Enterobacteriaceae transmission in our ICU setting, indicating effective infection control processes. We show that rapid molecular assays, such as our multiplex PCR, can detect common β-lactamase genes from bacterial culture within 40 minutes. These early molecular results would permit more informed selection of antibiotic therapy up to 1-2 days prior to 3GC-R confirmation with traditional culture methodologies. Further research is required to determine whether these molecular assays can be used directly on clinical specimens, and whether the implementation of molecular assays and more advanced technologies (e.g. WGS) is associated with improved patient outcomes in the hospital setting.

## Acknowledgements and funding

This study was funded by the Wishlist Sunshine Coast Health Foundation (awarded to AGS, KC, SS, and DSS). DSS and EPP were funded by Advance Queensland (grants AQRF13016-17RD2 and AQIRF0362018, respectively). We thank Dr Delaney Burnard (University of Queensland Centre for Clinical Research) and the Sunshine Coast University Hospital pathology laboratory for laboratory assistance, and Rhys T. White (University of Queensland) for helpful feedback on this manuscript.

## Transparency declarations

Dr. Harris reports research grants from Shionogi, Sandoz, and MSD and speaker’s fees from Pfizer, outside the submitted work. No conflicts are declared for remaining authors

